# MetaAMI: A Novel Meta-Learning Approach for Predicting In-Hospital Mortality in Acute Myocardial Infarction

**DOI:** 10.64898/2026.07.28.741291

**Authors:** Bulidierxin Tuerhanbayi, Xiaoting Fan, Jieqiong Wang, Shibiao Wan

## Abstract

Acute myocardial infarction (AMI) is one of the leading cardiovascular diseases worldwide and remains a major cause of mortality. Early risk prediction can help clinicians identify high risk patients shortly after admission and support timely monitoring and individualized treatment. Previous AMI risk assessment approaches predominantly rely on a single model structure or fixed feature representation, which may limit their ability to capture diverse risk related patterns and reduce predictive performance. To address these challenges, we propose MetaAMI, a random projection based meta-learning framework for AMI outcome prediction. Specifically, patients were selected based on ICD-9 and ICD-10 diagnostic codes for AMI from the Medical Information Mart for Intensive Care IV (MIMIC-IV) v3.1 database. Features were transformed through multiple random projections, with each random projection generating a distinct lower dimensional feature representation. Subsequently, baseline classifiers were trained on each lower dimension representation to predict in-hospital mortality among AMI patients. The predictions were aggregated to construct an integrated feature representation, which was used as input to a meta-learner architecture. By effectively integrating complementary information from diverse baseline models, the meta-learner refined the decision boundary and enhanced overall predictive performance. Survival analysis and SHapley Additive exPlanations (SHAP) analysis were further performed to evaluate clinical utility and interpret the model predictions. Benchmarking results based on MIMIC-IV dataset suggested that our MetaAMI consistently outperformed all the baseline classifiers across seven evaluation metrics including Accuracy, Area Under the Curve (AUC), F1 Score, G-Measure, Jaccard Index, Youden J, and Matthews Correlation Coefficient (MCC). In addition, feature importance analysis showed clinically relevant predictors of in-hospital mortality. In summary, MetaAMI provides an effective and robust solution for machine learning based AMI risk prediction. We anticipate that the application of MetaAMI will have a positive impact on clinical risk stratification and personalized treatment strategies for AMI.

## Introduction

Cardiovascular diseases are the leading cause of death in the European Union and the United States, accounting for approximately 30% of all fatalities [1]. Among them, acute myocardial infarction (AMI) is one of the most severe cardiovascular conditions and remains the primary cause of mortality and hospitalization worldwide [2, 3]. The in-hospital mortality rate of AMI, which reflects the quality of healthcare delivery and the efficacy of clinical interventions, varies substantially across different countries. The European Society of Cardiology [4] suggested that the mortality risk of AMI is modulated by numerous risk factors with strong predictive power for death. In addition to age, comorbidities, and elevated heart rate, abnormal alterations in several laboratory test indicators are also closely associated with this risk. However, conventional risk scores, such as TIMI [5], GRACE [6], and ACTION-GWTG [7] usually rely on a limited number of predefined variables, which may not fully reflect the complex physiological status of AMI patients. Therefore, more flexible approaches are needed to integrate diverse clinical information for accurate mortality risk prediction.

In the digital health era characterized by accessible and applicable massive datasets, artificial intelligence (AI) and machine learning (ML) algorithms can provide robust support for clinical decision making [8, 9]. The etiologies of AMI are intricate and its manifestations are highly heterogeneous; notably, the occurrence and progression of AMI are shaped by the combined influence of genetic, environmental, and behavioral factors [10]. Consequently, there is an increasingly urgent demand for the integrated analysis of multi-source data, including administrative data, laboratory test data and imaging data, to facilitate disease interpretation, clinical diagnosis and therapeutic decision making [11, 12]. The timely and accurate assessment and prediction of mortality risk constitute a critical component of clinical decision making, and machine learning algorithms hold substantial potential to improve the accuracy of such predictions.

In recent years, various AI/ML methods, including traditional machine learning models such as random forest, support vector machine and XGBoost [13, 14], as well as advanced deep learning models based on multilayer perceoptrons and Transformer architechtures[15], have been applied to AMI risk prediction, however, many approaches rely on a single model structure or fixed feature representation, limiting their ability to fully utilize the complementary advantages of different models. Moreover, although some models perform well on specific datasets, their stability and generalizability in different cohorts or real-world clinical settings still need to be verified.

To address these challenges, we introduce MetaAMI, a stacking based meta-learning framework that combines multiple random projection (RP) based feature representations and ensemble learning to predict in-hospital mortality risk among ICU AMI patients. RP effectively reduces feature dimensionality while preserving the geometric structure of the data [16]. Based on this, multiple baseline classifiers are trained under different RP representations, and their prediction results are further integrated for meta-learning to leverage complementary information across models [17]. In this way, the meta-learner can alleviate the systematic bias of a single model and improve the overall stability and robustness of the prediction.

## Method

### Cohort selection

This study utilized data from the Medical Information Mart for Intensive Care IV (MIMIC-IV) v3.1 database. MIMIC-IV is a large, publicly available, deidentified dataset comprising electronic health records of patients admitted to the emergency department or intensive care unit at Beth Israel Deaconess Medical Center in Boston. The database covers admissions from 2008 to 2019 and contains data for over 65,000 patients admitted to an ICU [18]. The database contains rich clinical characteristics, laboratory test results, vital signs, and complete medical records, providing high quality data support for this study. Patients were selected based on International Classification of Diseases, Ninth and Tenth Revision (ICD-9 and ICD-10) diagnostic codes for AMI (**Supplementary Table 1**). Patients meeting the following criteria were excluded from the study: (1) age less than 18 years at the time of first ICU admission; and (2) multiple ICU admissions. After these preprocessing steps, a total of 5854 patients were included in this study (**Fig. 1A**).

**Fig. 1.**
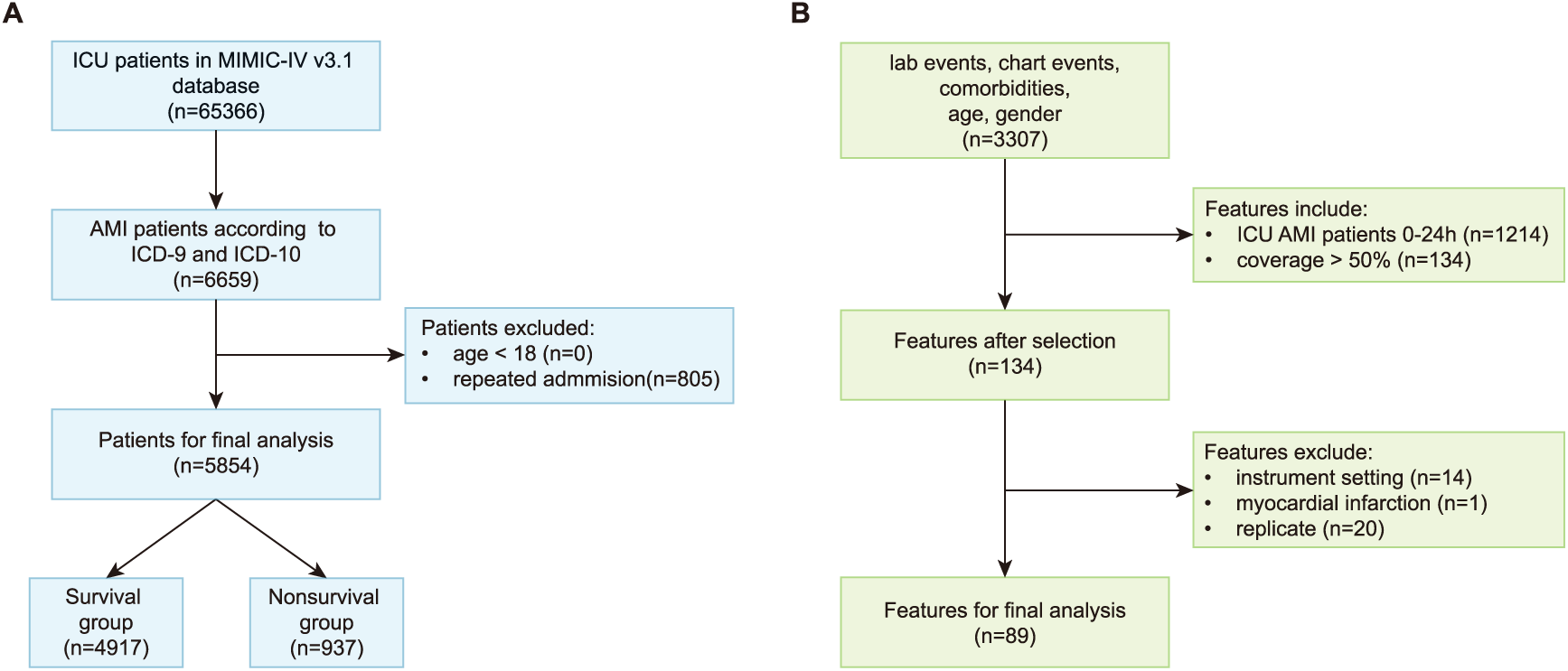
Flowchart of patient enrollment and feature filtering. **(A)** Flowchart of patient selection for ICU admissions with AMI from the MIMIC-IV database using ICD diagnosis codes. **(B)** Feature inclusion and exclusion criteria and preprocessing workflow.

### Features selection

To construct the input feature matrix for MetaAMI, we extracted four categories of variables from MIMIC-IV, including demographic characteristics, comorbidities, laboratory tests, and vital signs. The final input feature matrix consisted of 89 variables. Demographic characteristics included age and gender. Comorbidities were derived from the Charlson comorbidity table, and the Charlson Comorbidity Index (CCI) was also included as a quantitative measure of overall comorbidity burden. In addition, laboratory test results and vital signs within the first 24 hours after ICU admission were extracted from the MIMIC-IV database through an unsupervised feature selection way. Only features with a missing rate below 50% were retained. Instrument setting variables, myocardial infarction, and duplicated features were further removed (**Fig. 1B**). A total of 89 features were ultimately included, and the descriptive statistics of these features are shown in **Table 1**. To reduce the impact of missing data on model training, continuous variables were imputed using the median, and binary variables were imputed using the mode. Before model training, features were standardized using Z-score normalization. The primary outcome of this study was in-hospital mortality, defined based on the patient’s survival status at hospital discharge (survivor or non-survivor).

**Table. 1.**
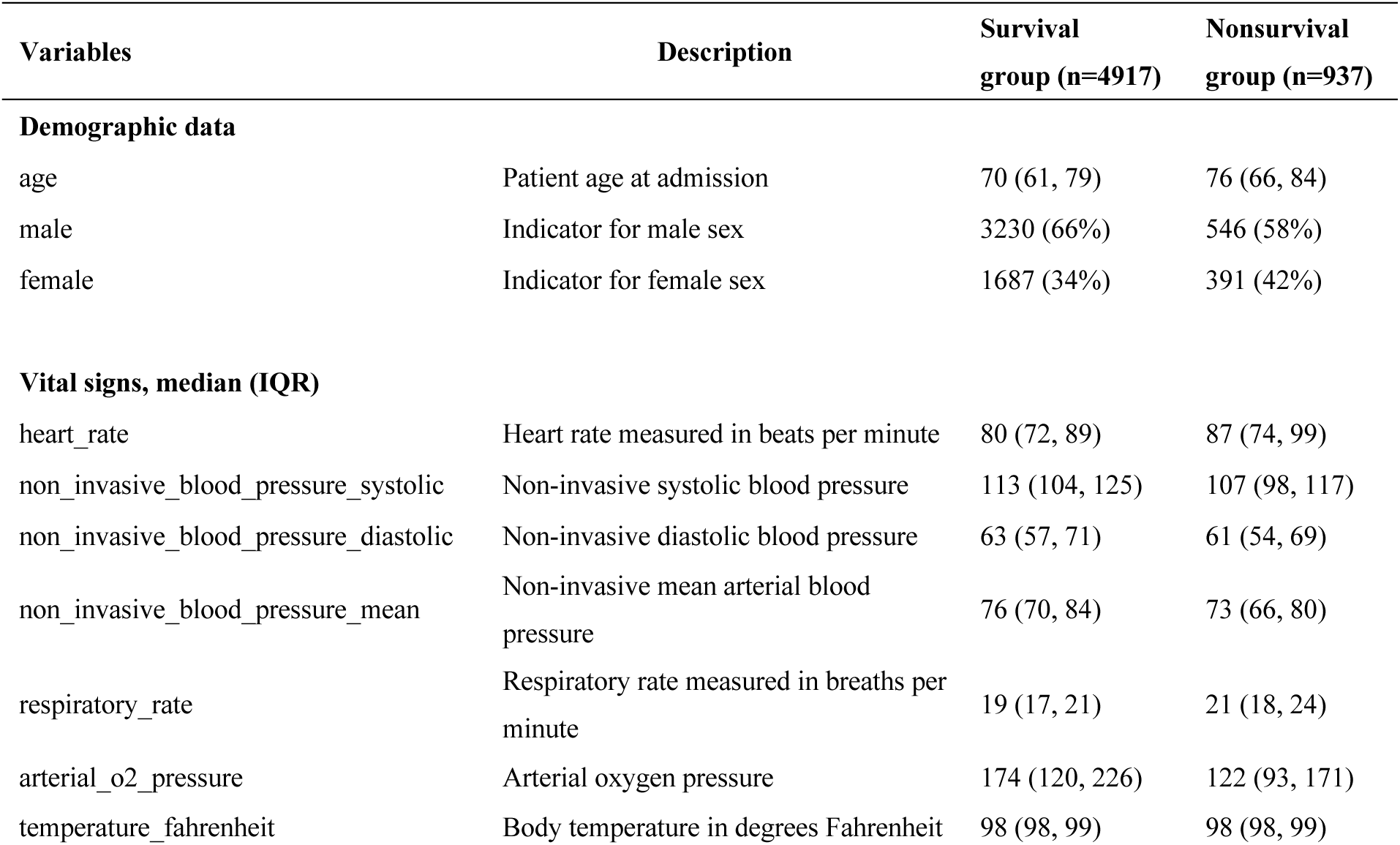
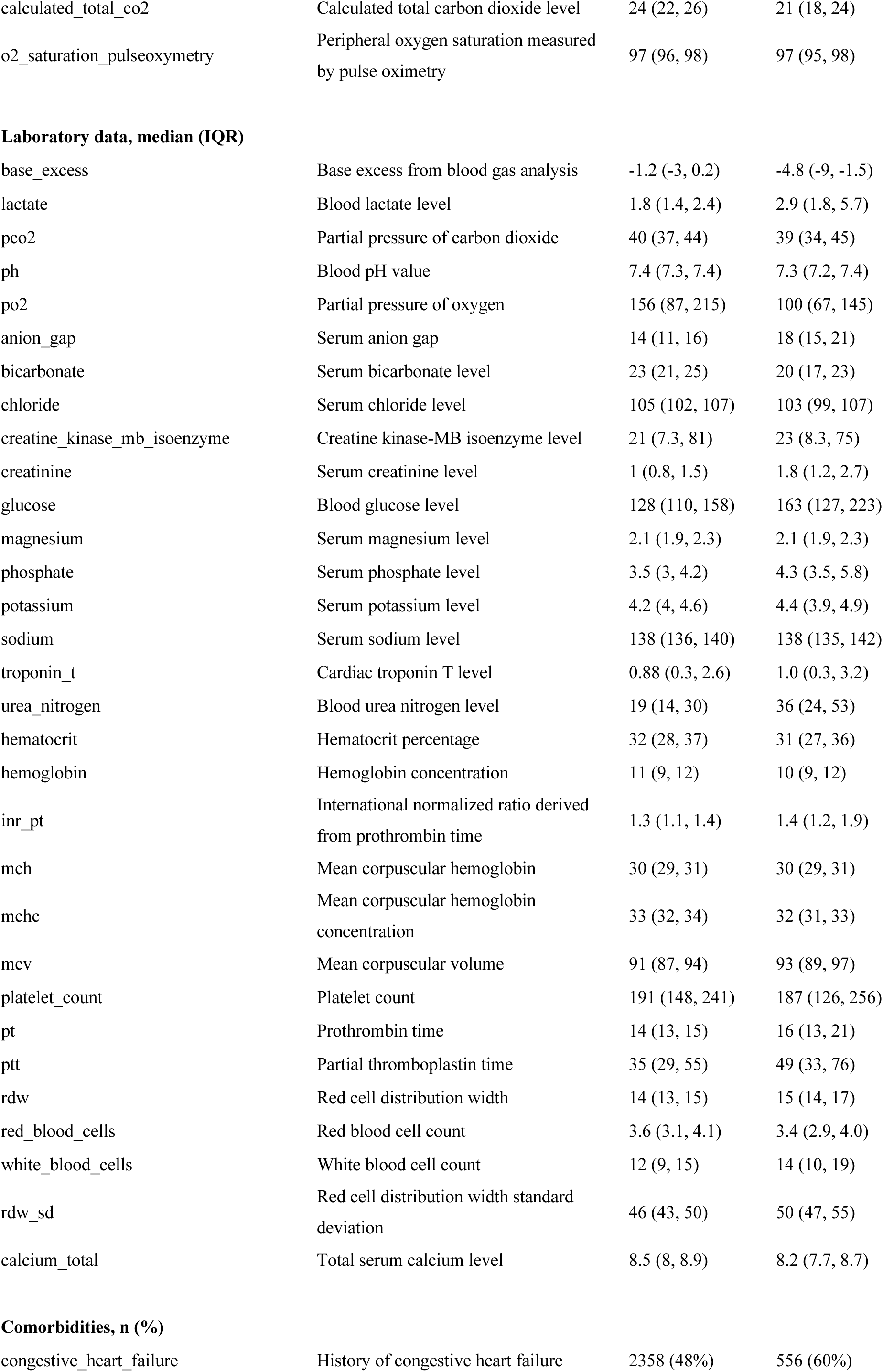
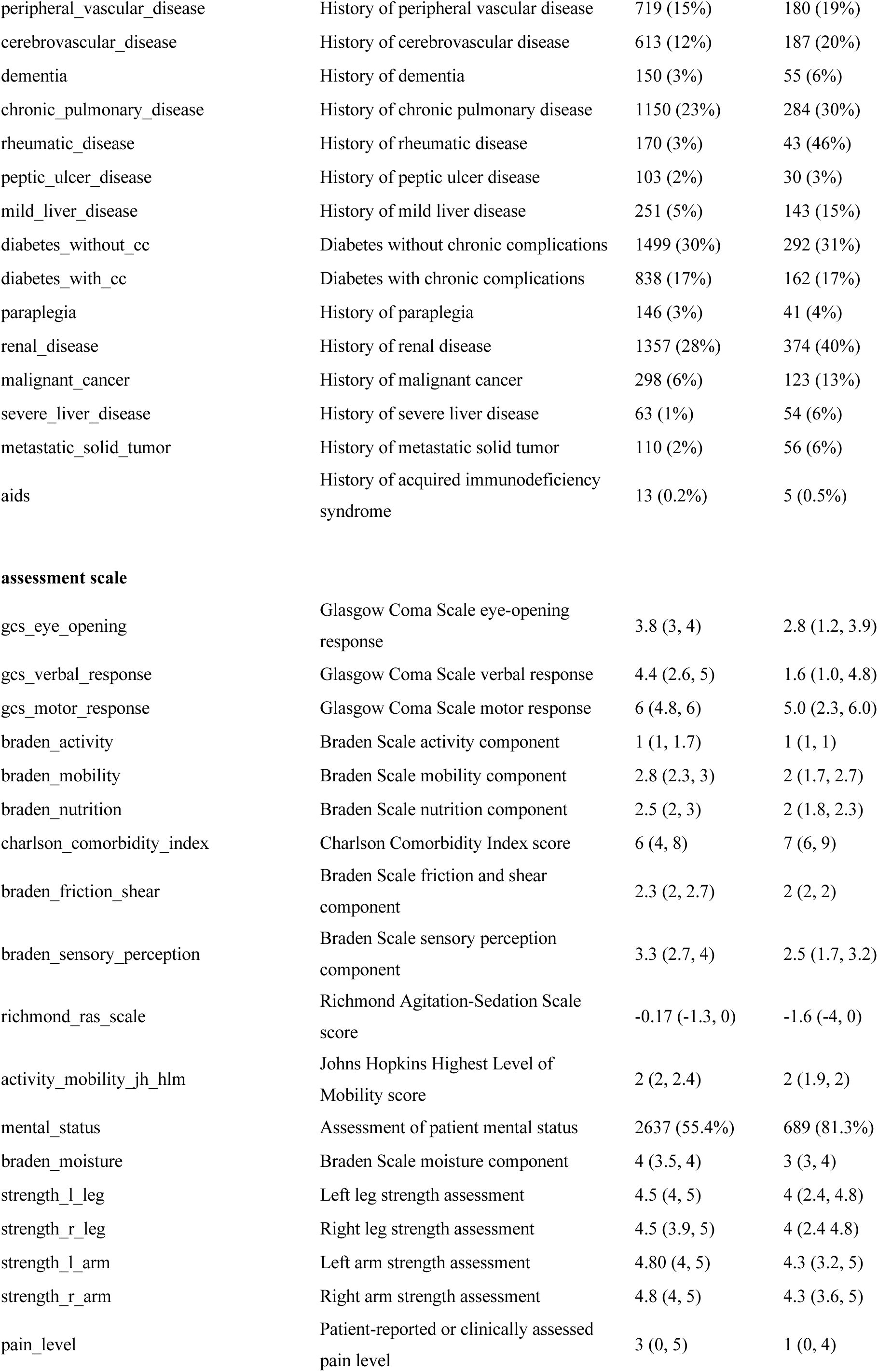
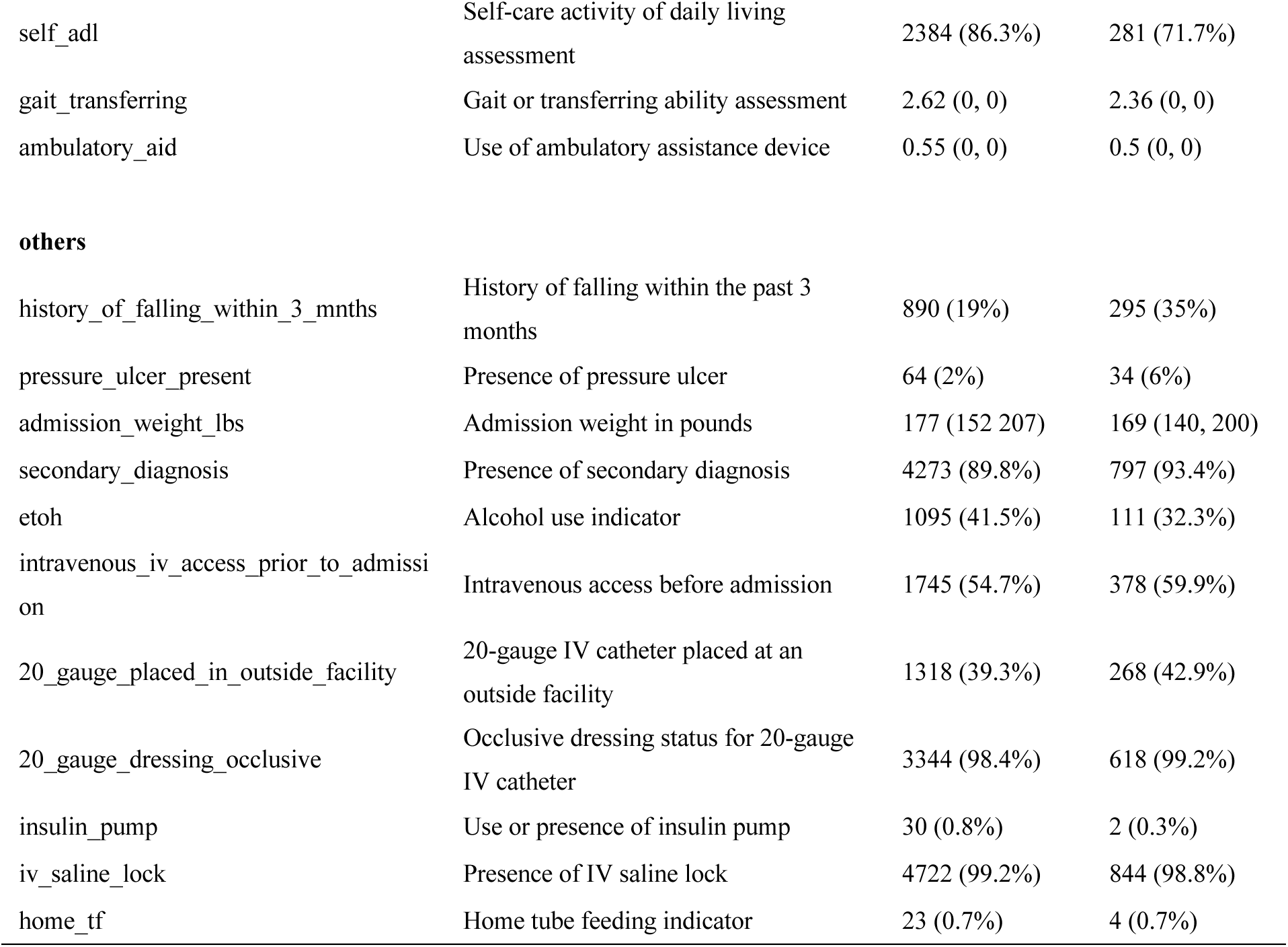
Descriptive statistics of the features in the survival and non-survival groups and their associations with in-hospital mortality. Continuous variables are presented as median (interquartile range), and categorical variables are presented as number (percentage). P values were calculated to compare the two groups. A P value < 0.05 was considered statistically significant.

### MetaAMI

In this study, the proposed predictive framework MetaAMI consists of three major steps: RP-based dimensionality reduction, baseline prediction, and stacking ensemble learning. The first step aimed to reduce feature dimensionality while preserving the overall structure of the original data. Specifically, the original feature data were subjected to multiple times of dimensionality reduction using the RP method. Compared with PCA, RP is computationally less expensive and can approximately preserve pairwise distances between samples after dimensionality reduction [19, 20]. Multiple independent RPs further introduced diversity in the resulting low dimensional representations, as each projection was generated using a different random transformation matrix. Second, each RP transformed feature matrix was used to train a set of baseline classifiers and generate baseline predictions. Third, these prediction outputs were integrated by a stacking meta-learner to produce the final prediction of in-hospital mortality risk.

The baseline classifiers included Logistic Regression (LR) [21], Random Forest (RF) [22], Extreme Gradient Boosting (XGBoost) [23], Support Vector Machine (SVM) [24], Multilayer Perceptron (MLP) [25], Autoencoder-Multilayer Perceptron (AE-MLP) [26], Optimal Transport-Multilayer Perceptron (OT-MLP) [27], and Transformer [28].

In the meta-learning stage, base classifiers were further integrated. Specifically, based on the performance ranking of the base classifiers on the validation set, we evaluated three joint stacking settings: the top 4, top 6, and top 8 performing base classifiers. For each setting, the predicted probabilities of the selected models were concatenated to form the input features of the meta-learner. In addition, all pairwise combinations within the top 4 classifier set were further evaluated. Pairwise analysis was restricted to the top 4 classifiers to limit the number of model combinations. This strategy enables the meta-learner to systematically integrate predictive information from different random projection spaces and different model architectures.

The stacking based meta-learning model generated the final binary probability predictions by integrating complementary information from different base classifiers. To determine the optimal meta-learning strategy, we systematically evaluated six candidate meta-classifiers, including SVM, LR, RF, XGBoost, MLP, and Transformer, using the area under the receiver operating characteristic curve (AUC) as the primary performance evaluation metric. The model with the best performance was ultimately selected as the final meta-learning framework. In this study, SVM achieved the best performance and was selected as the final meta-classifier. Detailed hyperparameter settings for all classifiers are provided in (**Supplementary Table 2**).

Model training and evaluation were performed using ten times ten-fold cross-validation. Model performance was comprehensively evaluated using seven different metrics, including AUC, accuracy, F1 score, Youden index, MCC, G-measure, and Jaccard index. Youden index, defined as sensitivity + specificity - 1, was used to determine the optimal classification threshold [29]. MCC incorporated all four elements of the confusion matrix and provided a balanced performance measure for binary classifications. G-measure evaluated the geometric mean of sensitivity and specificity, and the Jaccard index quantified the overlap between predicted and actual death cases.

## Statistical Analysis

Continuous variables are presented as median and interquartile range (IQR). For normally distributed continuous variables, group comparisons were performed using a two-sided Student’s t-test; for non-normally distributed continuous variables, the Wilcoxon Mann-Whitney U test was applied. Categorical variables were presented as counts and percentages, and group comparisons were performed using the chi-square test or Fisher’s exact test. Statistical differences between model classification performances were evaluated using the McNemar test to compare prediction outcomes on the same patient cohort [30]. A *p* value < 0.05 was considered statistically significant for all statistical tests.

### Kaplan-Meier Survival Curve Analysis

The in-hospital mortality prediction probability for each patient was calculated using the meta-learner. The optimal threshold was determined using the Youden index. Patients were divided into low risk, medium risk, and high risk groups based on risk score. Kaplan-Meier (KM) survival curves were constructed using the survival package in R, with ICU admission time as the starting point and in-hospital death or discharge as the endpoint. The log-rank test was used to compare survival differences between the three groups, with a *p* value < 0.05 considered statistically significant. The time-to-event analysis was performed to evaluate the model’s ability to stratify mortality risk over time and to validate its clinical application value in early prognostic evaluation of critically ill AMI patients.

### SHAP Analysis

To evaluate the importance of individual feature in model predictions, a feature ranking method was performed [31]. Shapley values derived from cooperative game theory were used to quantify the relative contribution of each input variable to the model output. To address the black box problem of machine learning models and enhance clinical interpretability, we applied the SHAP method to the best performing baseline classifier for explanatory analysis. A beeswarm plot of the top 25 most important features was generated to illustrate the relationship between feature values and their corresponding SHAP values, thereby revealing both the direction and magnitude of each feature’s influence on model predictions.

## Results

### Design of the MetaAMI

To accurately predict the in-hospital mortality in ICU patients with AMI, we proposed MetaAMI, a stacking based meta-learning framework that combines random projection with ensemble learning (**Fig. 2**). First, we applied random projection multiple times to perform dimensionality reduction, aiming to extract clinically meaningful latent information while increasing the variability of feature representations. The features processed by random projection were then used as inputs to train multiple baseline classifiers independently. Next, the prediction outputs generated by these baseline classifiers were concatenated and fed into the meta-learner. By integrating complementary predictive information from different baseline models, the meta-learner further refined the classification decision boundary and produced the final mortality risk prediction.

**Fig. 2.**
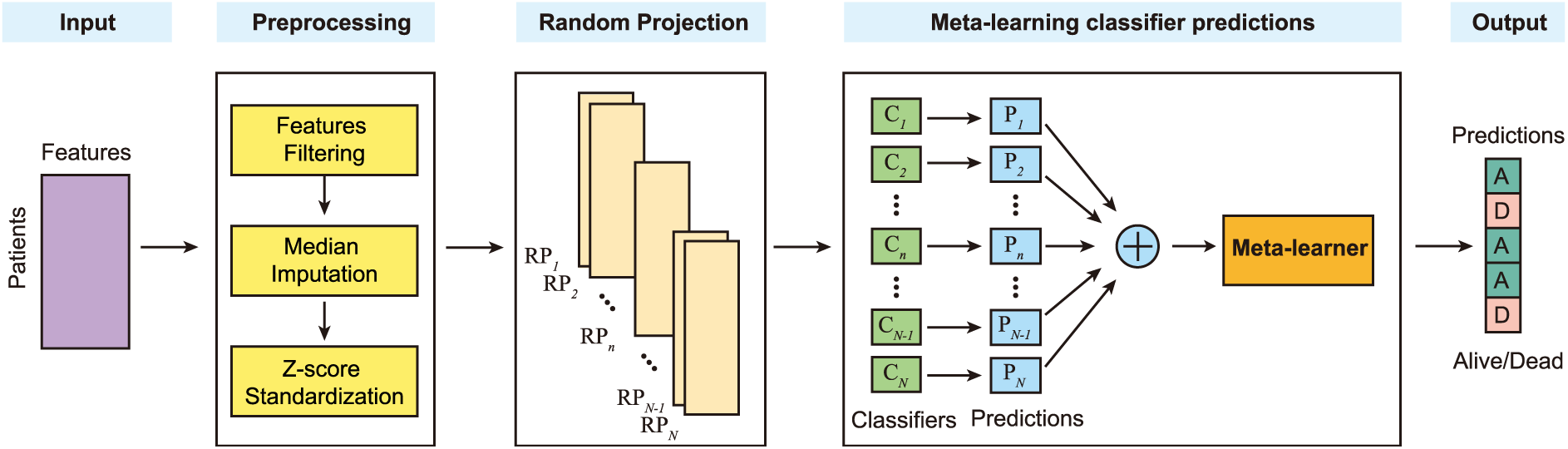
Design of MetaAMI. MetaAMI consists of four steps for AMI in-hospital mortality prediction: data preprocessing, multiple random projection based dimensionality reduction, baseline classifiers predictions, meta-learner prediction. RP: random projection, C: classifier, P: prediction, A: alive, D: dead.

To identify the optimal configuration of MetaAMI, we systematically evaluated its predictive performance across different classifier combinations and random projection settings. First, eight base classifiers were trained independently and ranked based on their predictive performance, serving as references for subsequent model selection. Next, we tested MetaAMI under different random projection dimension ranging from 20 to 50 to identify the optimal dimensionality (**Supplementary Fig. 1A**). The AUC continuously improved as the projection dimension increased from 20 to 40, and then decreased. Based on the optimal projection dimension of 40, we further evaluated different combinations of baseline models and observed that the combination of MLP and AE-MLP achieved the best performance (**Supplementary Fig. 1B**). Furthermore, we explored the effect of using multiple independent RP matrices on model performance (**Supplementary Fig. 1C**). The AUC steadily improved as the number of RPs increased to approximately 30. Based on the optimal base classifier combination identified above (MLP and AE-MLP), we further compared the predictive performance of different meta-learners under the same RP settings. SVM demonstrated the best performance, yielding the final optimal meta-learner architecture (**Supplementary Fig. 1D**).

### MetaAMI Outperformed Individual Baseline Classifier for AMI In-Hospital Mortality Prediction

Based on the optimal structure, we compared the predictive performance of MetaAMI with individual baseline classifier. MetaAMI achieved the best performance across all evaluation metrics, including AUC, accuracy, F1 score, youden index, MCC, G-measure, and jaccard index. Specifically, accuracy improved by 2.4% (**Fig. 3A**), while AUC increased by 0.8% (**Fig. 3B**) and the F1 score improved by 2.5% (**Fig. 3C**). The G-measure increased by 1% (**Fig. 3D**). MetaAMI also achieved improvements of 2.5% in Jaccard index (**Fig. 3E**), 1.8% in Youden index (**Fig. 3F**), and 2.7% in MCC (**Fig. 3G**). The confusion matrices further showed that MetaAMI provided the most balanced classification performance, correctly identifying 82% of survival cases and 83% of death cases (**Fig. 4A**). AE-MLP correctly classified 75% of survival cases and 88% of death cases (**Fig. 4B**), while MLP correctly classified 76% and 87%, respectively (**Fig. 4C**). OT-MLP correctly identified 79% of survival cases and 85% of death cases (**Fig. 4D**). The Transformer correctly classified 77% of survival cases and 87% of death cases (**Fig. 4E**), whereas SVM correctly classified 78% and 83%, respectively (**Fig. 4F**). LR, RF, and XGBoost, however, showed a strong tendency to predict death, achieving death case classification rates of 98%, 99%, and 99%, respectively, but substantially lower survival case classification rates of 42%, 29%, and 32% (**Fig. 4G-I**). In addition, McNemar’s tests demonstrated that MetaAMI achieved statistically significantly better classification performance than all individual baseline classifiers (*p* < 0.0001) (**Supplementary Table. 3**). Overall, these results highlight the advantages of the proposed meta-learning framework: by integrating complementary information from multiple base classifiers, MetaAMI yields more stable and accurate predictions than any individual model alone.

**Fig. 3.**
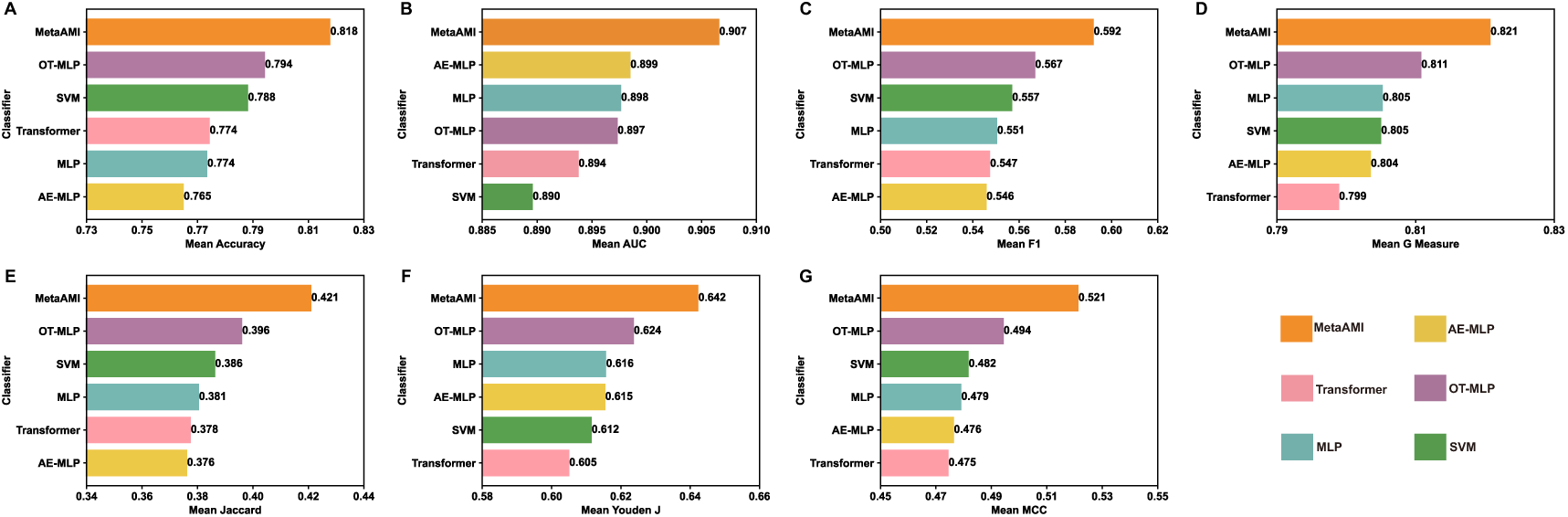
Performance evaluation of MetaAMI. MetaAMI outperformed the baseline models across multiple evaluation metrics, including **(A)** Accuracy, **(B)** Area Under the Curve (AUC),**(C)** F1 Score, **(D)** G-Measure, **(E)** Jaccard Index, **(F)** Youden J, and **(G)** Matthews Correlation Coefficient (MCC).

**Fig. 4.**
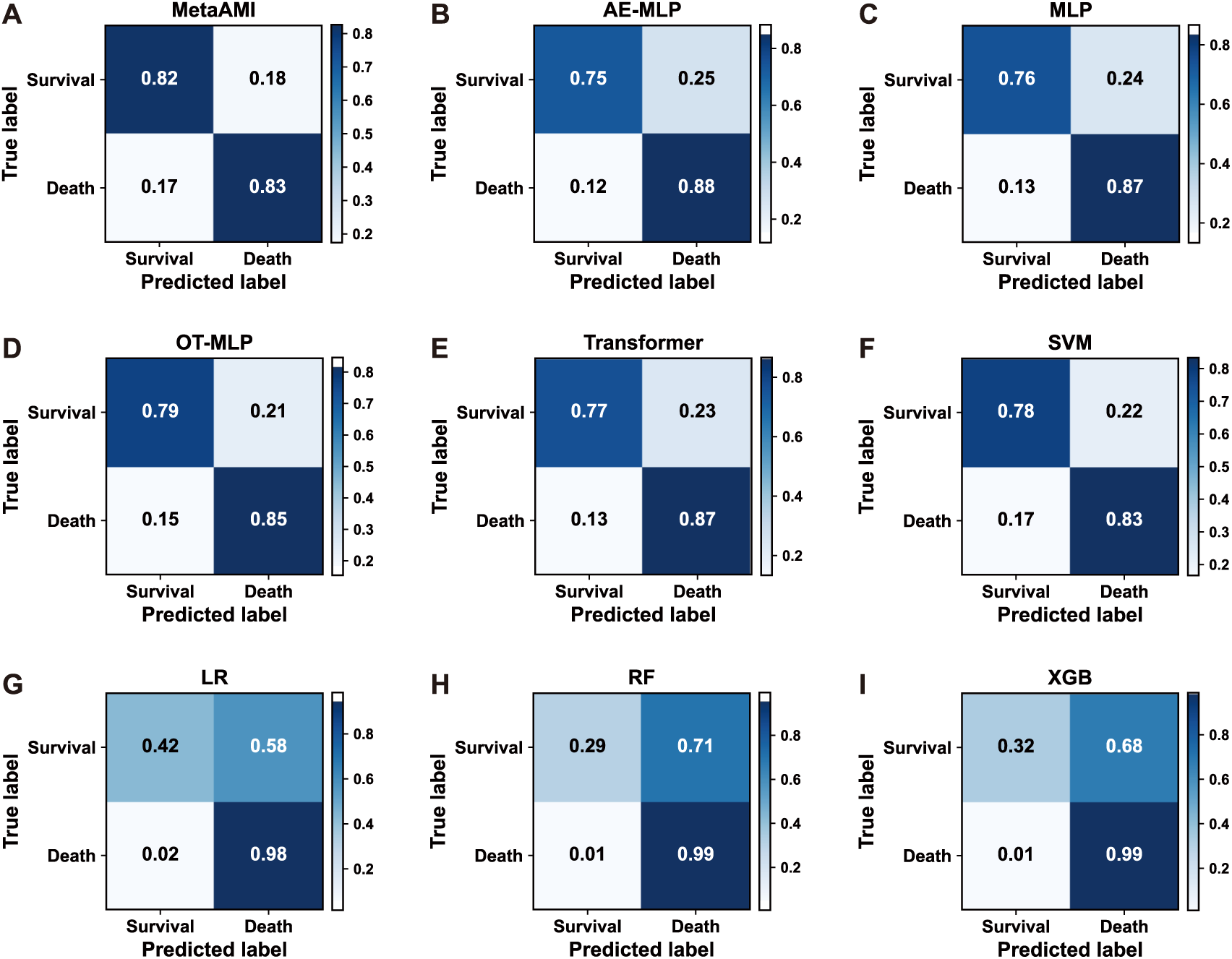
Confusion matrices of the evaluated models. Rows represent the true labels and columns represent the predicted labels. Each matrix shows the proportions of true positives, true negatives, false positives, and false negatives. The figure shows the confusion matrices for (**A**) MetaAMI, (**B**) AE-MLP, (**C**) MLP, (**D**) OT-MLP, (**E**) Transformer, (**F**) SVM, (**G**) LR, (**H**) RF, and (**I**) XGBoost

### MetaAMI Outperformed State-of-the-Art Approaches for AMI In-Hospital Mortality Prediction

We further compared MetaAMI with three state-of-the-art appraoches, including myocardial infarction graph neural network (MI-GNN)[32], conditional medical generative adversarial network (cMedGAN)[33], and Acute Physiology Score III (APS III)[34]. MI-GNN is a graph neural network based prediction method that first constructs a patient similarity graph to represent relationships among patients, and then applies a Graph Transformer to learn graph based patient representations for in-hospital mortality prediction. cMedGAN is a generative data augmentation based prediction method, which first uses a conditional medGAN model to generate synthetic patient samples in the latent space and then combines these synthetic samples with real training samples to train an MLP classifier. APS III is a traditional clinical severity scoring system, and in this study, its predicted probability was used to evaluate the risk of in-hospital mortality in patients with AMI. The results showed that, compared with MI-GNN, cMedGAN, and APS III, MetaAMI achieved better performance across multiple evaluation metrics, including AUC, accuracy, F1 score, Youden index, MCC, G-measure, and Jaccard index. Specifically, accuracy improved by 2.1% (**Fig. 5A**), while AUC increased by 2% (**Fig. 5B**) and the F1 score improved by 3% (**Fig. 5C**). The G-measure increased by 1.9% (**Fig. 5D**). MetaAMI also achieved improvements of 2.9% in Jaccard index (**Fig. 5E**), 3.6% in Youden index (**Fig. 5F**), and 3.6% in MCC (**Fig. 5G**).

**Fig. 5.**
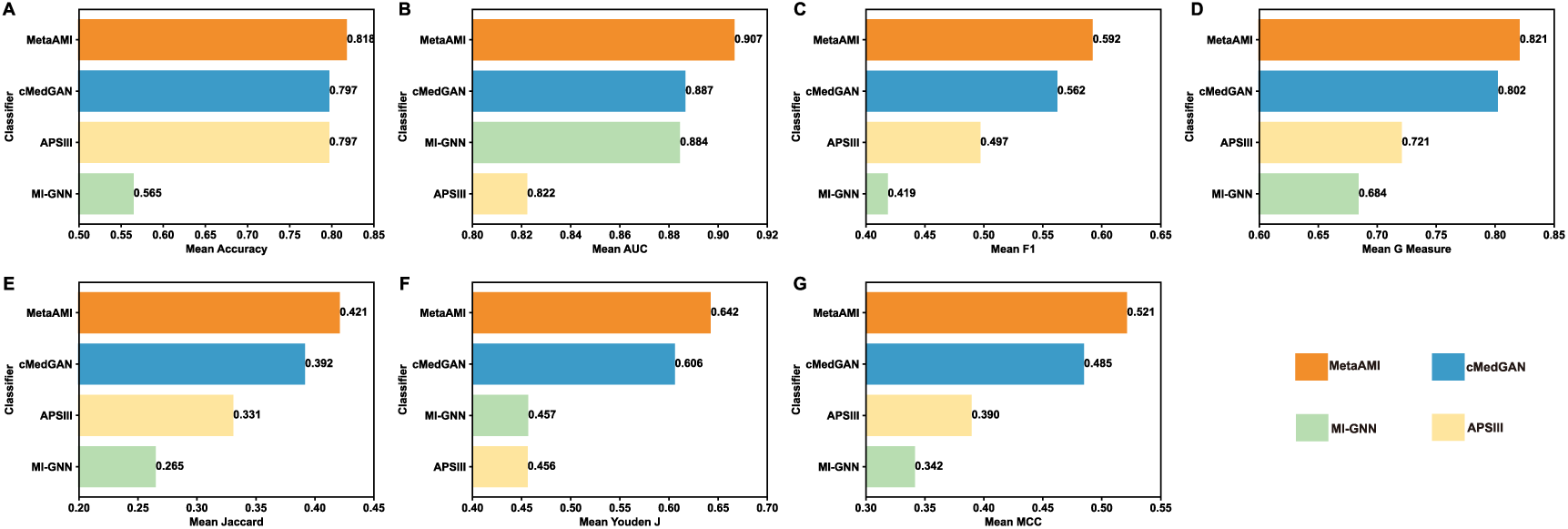
Benchmark performance evaluation of MetaAMI. MetaAMI showed improved performance compared with the benchmark models across multiple evaluation metrics, including **(A)** Accuracy, **(B)** AUC, **(C)** F1 Score, **(D)** G-Measure, **(E)** Jaccard Index, **(F)** Youden J, and **(G)** MCC.

### MetaAMI Enables Effective Risk Stratification and Provides Clinical Benefit

To systematically evaluate the potential clinical utility of MetaAMI in the ICU setting, we analyzed its performance from two aspects: risk stratification ability and clinical net benefit. Based on the predicted risk scores, patients were divided into three risk groups using the 50^th^ and 90^th^ percentiles as cutoffs: low risk, medium risk, and high risk groups. KM survival curves were plotted to compare survival outcomes among the three groups (**Fig. 6A**). The result showed a statistically significant difference in survival across the three groups (log-rank test, *p* < 0.0001), indicating that MetaAMI can effectively distinguish ICU admitted AMI patients with different mortality risks.

**Fig. 6.**
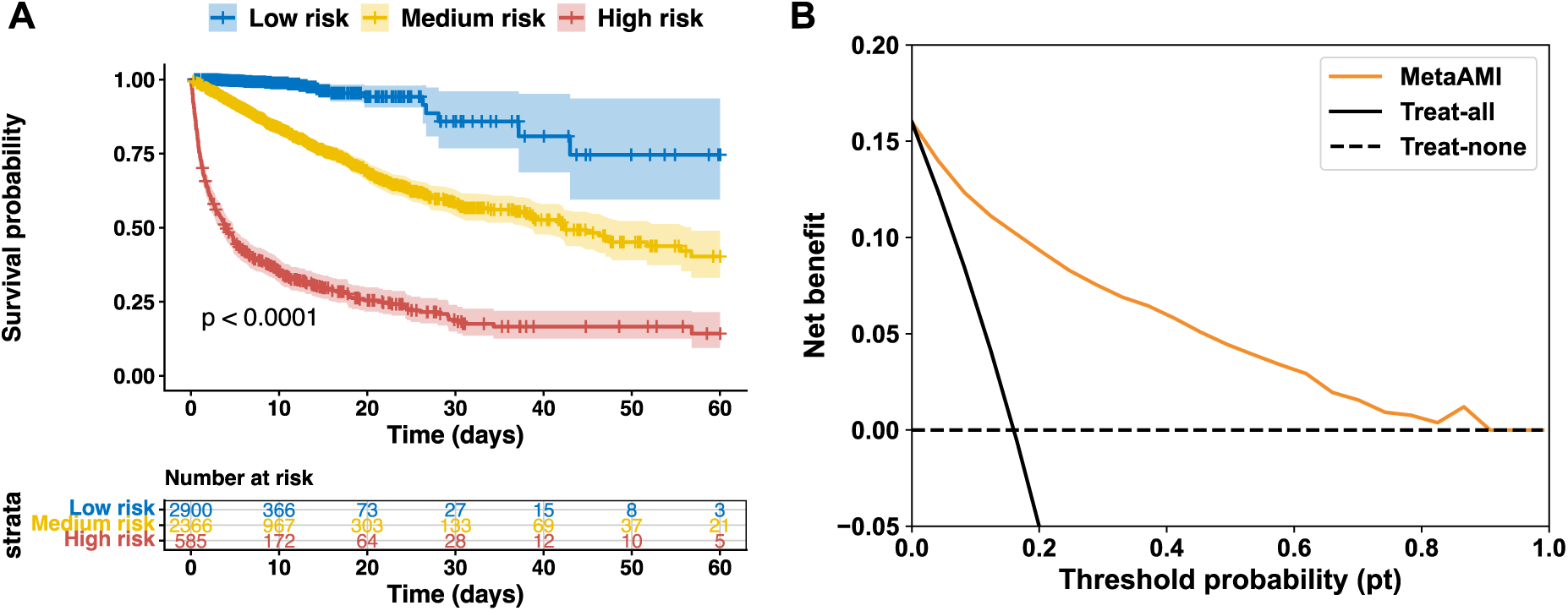
Clinical utility of MetaAMI. **(A)** Kaplan-Meier survival curves and risk table for low risk, medium risk, and high risk groups identified by MetaAMI. **(B)** Decision curve of MetaAMI.

Furthermore, we used Decision Curve Analysis (DCA) to evaluate the clinical net benefit of MetaAMI across different threshold probabilities (pt). Within a range of 0 to 0.9, the net benefit curve of MetaAMI consistently remained above the treat-all and treat-none strategies (**Fig. 6B**). This finding suggests that risk stratification guided by MetaAMI yields superior clinical benefit compared with uniform intervention or no intervention. By accurately identifying high risk patients while reducing unnecessary misclassifications, MetaAMI achieves a favorable balance between improving intervention efficiency and minimizing the risk of overtreatment.

### Feature Importance Analysis Identified Key Predictors of AMI In-Hospital Mortality

Based on SHAP analysis of the MLP and AE-MLP model, we obtained the relative importance of the top 25 features and their contributions to the model output (**Fig. 7**). Among all features, Braden nutrition and Charlson Comorbidity Index (CCI) was evaluated as the most influential variable affecting model predictions, followed by Mean Corpuscular Hemoglobin Concentration (MCHC), urea nitrogen, and age. Higher CCI is generally associated with a higher risk of in-hospital death, while lower CCI tends to shift the model predictions toward a survival outcome. Similar interpretations apply to other key features, and the directionality of feature importance for most features is consistent with established clinical knowledge. Furthermore, most of the top 25 key features in the model showed statistically significant differences between the survival and non-survival groups (**Figure. 8**) and were highly consistent with the features selected by experts, with an overlap rate of 90% (**Supplementary Table 4**). These results further support the validity and clinical plausibility of the SHAP based interpretations and demonstrate that machine learning methods can effectively identify outcome relevant features from high dimensional clinical data.

**Fig. 7.**
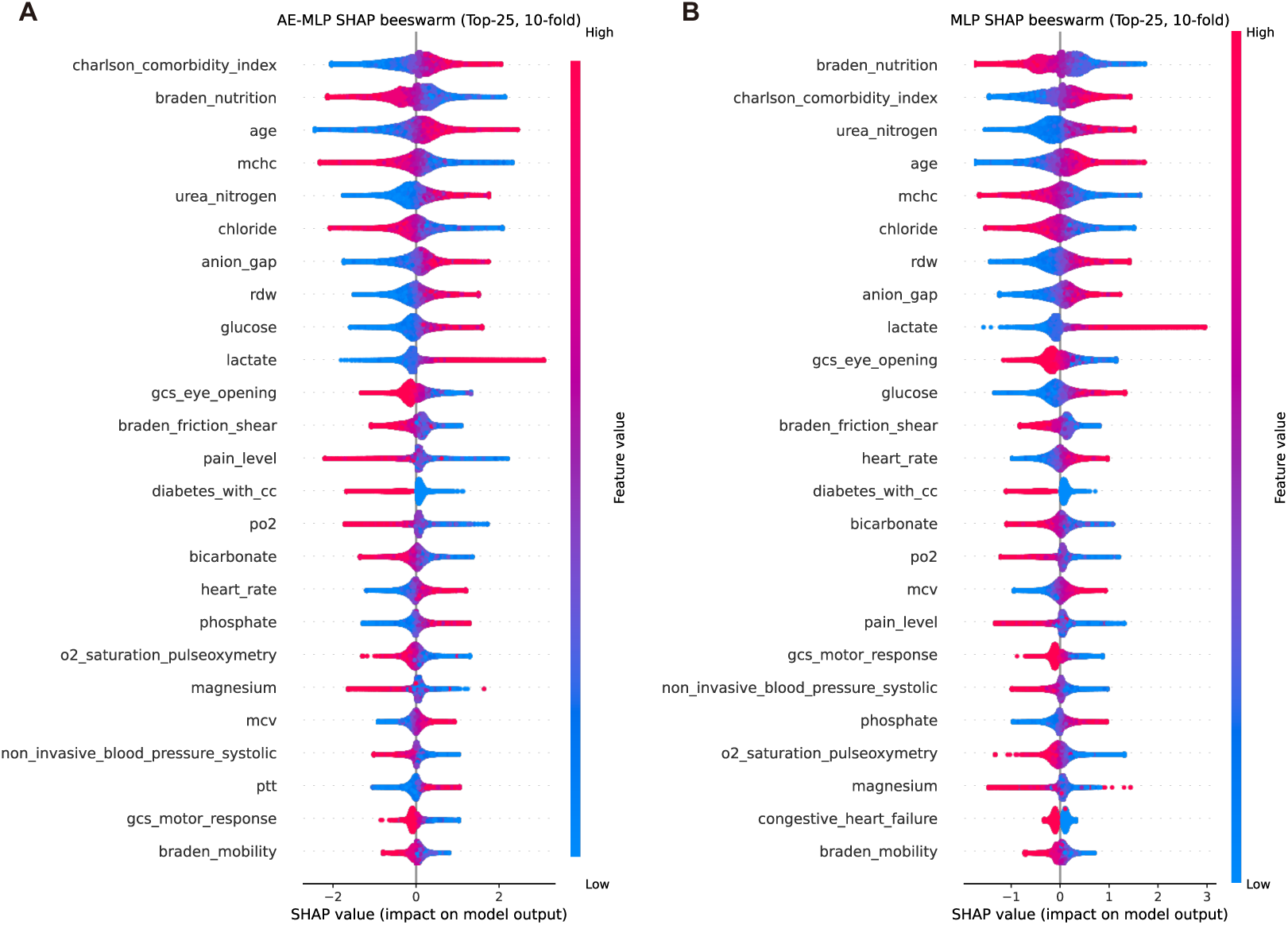
Impact of input features on model predictions. Top 25 features ranked by SHAP values for the MLP **(A)** and AE-MLP **(B)** models. Each dot represents an individual patient. The color indicates the magnitude of the feature value. A positive SHAP value indicates an increased predicted risk of mortality, while a negative SHAP value indicates a decreased predicted risk of mortality.

**Fig. 8.**
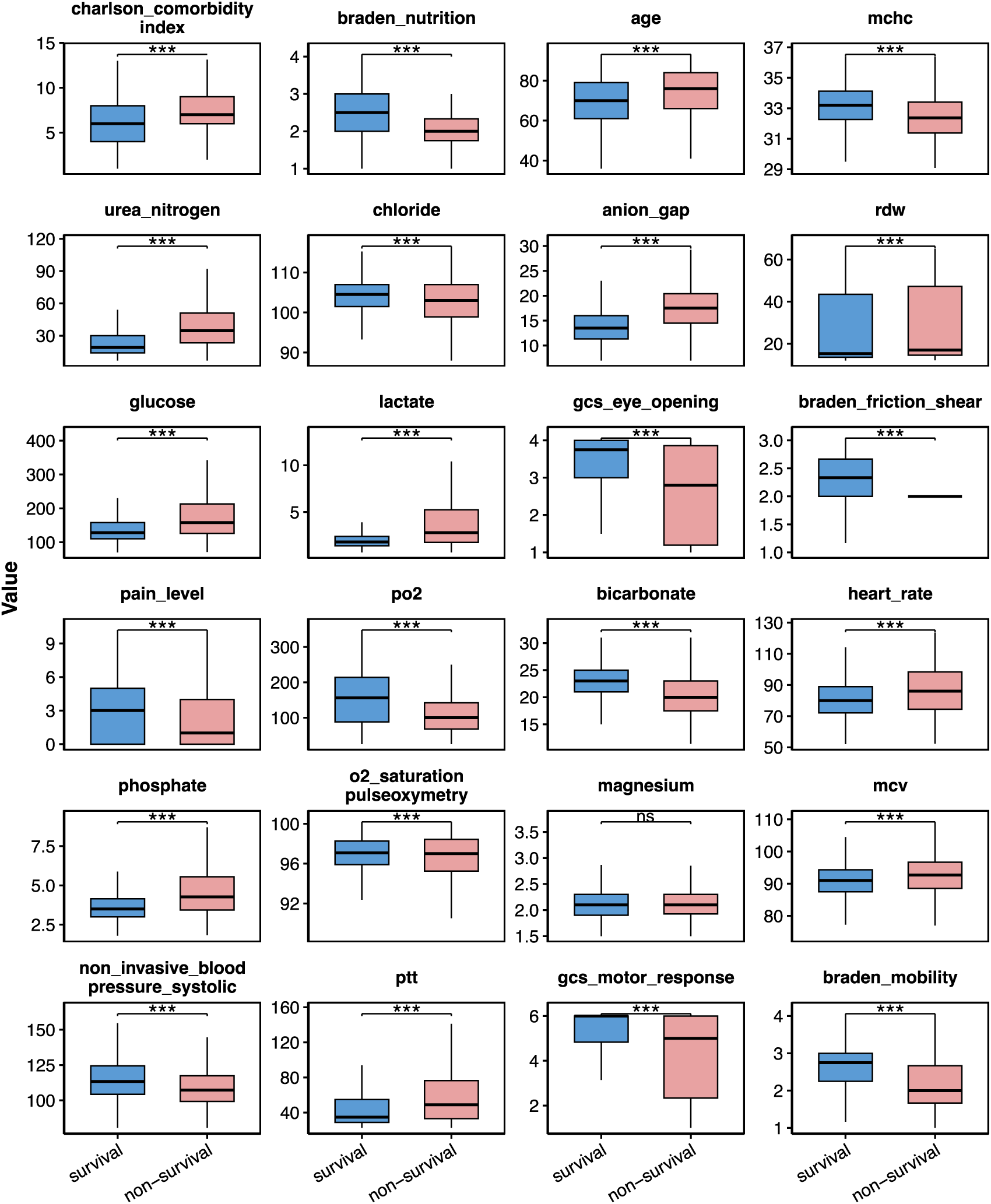
Boxplots of continuous SHAP selected features in the survival and non-survival groups. The distributions of the selected features are compared between the survival and non-survival groups. Boxes indicate the median and interquartile range, whiskers represent the range excluding outliers. *p* < 0.05*, *p* < 0.01**, *p* < 0.001***, ns: not significant.

## Discussion

In this study, we proposed a meta-learning framework called MetaAMI that integrated RP based dimensionality reduction with stacking ensemble learning to predict the in-hospital mortality of AMI patients in the ICU. The framework aggregated prediction outputs from multiple base classifiers trained on diverse RP embeddings, enabling the meta-learner to capture complex nonlinear relationships in routine clinical data and thereby enhancing model robustness and predictive stability. Our results showed that the optimal MetaAMI achieved an improvement of approximately 0.8% in AUC compared with the best individual base classifier. Across multiple evaluation metrics, MetaAMI consistently outperformed base classifiers. For ICU AMI patients, effective risk prediction requires not only accurate identification of high risk patients but also avoidance of excessive false positive predictions that may lead to unnecessary interventions. MetaAMI demonstrated a good balance between sensitivity and specificity, reflecting its potential applicability in real clinical scenarios.

Kaplan-Meier survival analysis based on MetaAMI’s predictions showed significant survival differences among the low risk, medium risk, and high risk groups. The finding indicated that routinely collected clinical data had the ability to distinguish ICU AMI patients across different risk groups. Further decision curve analysis suggested that risk stratification guided by MetaAMI provided greater net clinical benefit than strategies of intervening on all patients or not intervening at all. By accurately identifying truly high risk individuals while minimizing unnecessary misclassification, MetaAMI supported more efficient and targeted clinical decision making, which may facilitate timely interventions and potentially improve clinical outcomes.

We performed SHAP analysis for the interpretability of the prediction results and found that the influence of most features on the prediction was consistent with clinical practice and previous evidence. Braden scores (such as braden nutrition, braden friction shear, braden mobility) and GCS scores suggest that poor nutritional status, reduced mobility, and decreased level of consciousness are strongly associated with increased mortality risk [35, 36]. CCI and specific comorbidities such as congestive heart failure and diabetes were also identified as important predictors. Diabetes contributes to the development of coronary artery disease [37]. Patients with multiple comorbidities have a higher risk of death following AMI [38]. Hematological indicators such as MCHC, MCV, and RDW may reflect decreased blood oxygen carrying capacity and influence outcomes [39, 40]. Metabolic and electrolyte related indicators such as chloride, anion gap, bicarbonate, lactate, glucose, magnesium, and phosphorus reflect metabolic and acid base balance disorders, which are common in critically ill patients [41, 42]. Unstable Vital signs such as increased heart rate, decreased levels of blood pressure or impaired oxygenation usually indicate that the patient is in a critical condition and is closely related to a poor prognosis [43]. These highly important features collectively point to a systemic stress response under myocardial ischemia, including metabolic imbalance and diminished physiological reserve, ultimately contributing to increased mortality risk.

Moreover, except for magnesium, the top 25 key features identified by SHAP showed statistically significant differences between the survival and non-survival group, and were highly consistent with the key features selected by the expert, with an overlap rate of 90%. This high level of agreement suggests that machine learning methods can reliably capture clinically meaningful prognostic features from high dimensional data, demonstrating strong consistency with expert knowledge.

This study still has some limitations. First, the data were derived from the publicly available MIMIC-IV database, and the generalization ability of the model in other medical institutions or different populations needs to be confirmed through multicenter external validation. Second, this study mainly relied on structured clinical data and did not include other relevant modalities or longitudinal follow up data, which may limit the model’s ability to capture dynamic disease progression. Future work will focus on incorporating multimodal data to further enhance predictive performance and optimizing the meta-learning architecture to reduce computational complexity while maintaining robust performance.

## Conclusion

Overall, the proposed MetaAMI framework integrated random projection based dimensionality reduction with stacking ensemble learning. By integrating information from multiple random projections and various baseline models, MetaAMI achieved superior performance in predicting in-hospital mortality among ICU patients with AMI compared with individual baseline classifiers and benchmark models. MetaAMI effectively stratified patients into high risk, medium risk, and low risk categories. Furthermore, decision curve analysis showed that MetaAMI provided potential clinical net benefit across a specific threshold range, suggesting that it may help identify high risk patients early and assist clinicians in making more timely and targeted intervention strategies. SHAP analysis identified the key clinical features contributing to the model predictions and provided insight into how these features influenced mortality risk, thereby improving model interpretability. These results indicate that MetaAMI may serve as a clinical decision support tool for the early risk assessment and more personalized clinical management of critically ill AMI patients.

## Author Contribution

The idea for this study was conceived and designed by SW and XF. SW and BT developed the method. XF selected features. BT performed the experiment, and analyzed the data. All authors participated in the writing and revision of the paper. The manuscript was approved by all authors.

## Data Availability

All data used in this study are publicly available from MIMIC-IV v3.1 database.

## Code availability

All code used for MetaAMI is publicly available and can be found on GitHub at https://github.com/wan-mlab/MetaAMI.

