## Supplementary Figures and Tables for "MetaAMI: A Novel Meta-Learning Approach for Predicting In-Hospital Mortality in Acute Myocardial Infarction"

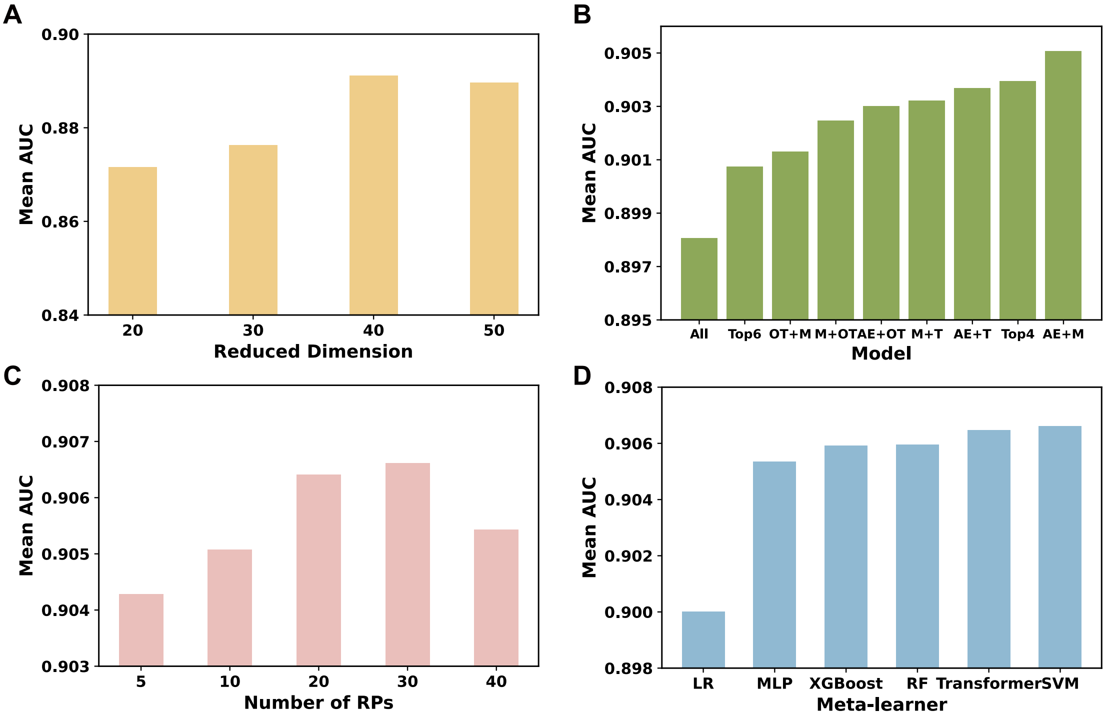


**Supplementary Fig. 1 Selection of RP and model configuration. (A)** Performance of MetaAMI across different RP dimensions. The best performance was achieved at 40 dimensions. **(B)** Performance of different baseline model combinations using 10 RPs. The combination of AE-MLP and MLP achieved the best performance. **(C)** Performance of the AE-MLP and MLP ensemble under different numbers of RPs. The optimal performance was observed when the number of RPs was 30. **(D)** Comparison of different meta-learners built on the AE-MLP and MLP ensemble, with SVM achieving the best performance. M: MLP; OT: OT-MLP; AE: AE-MLP; T: Transformer.

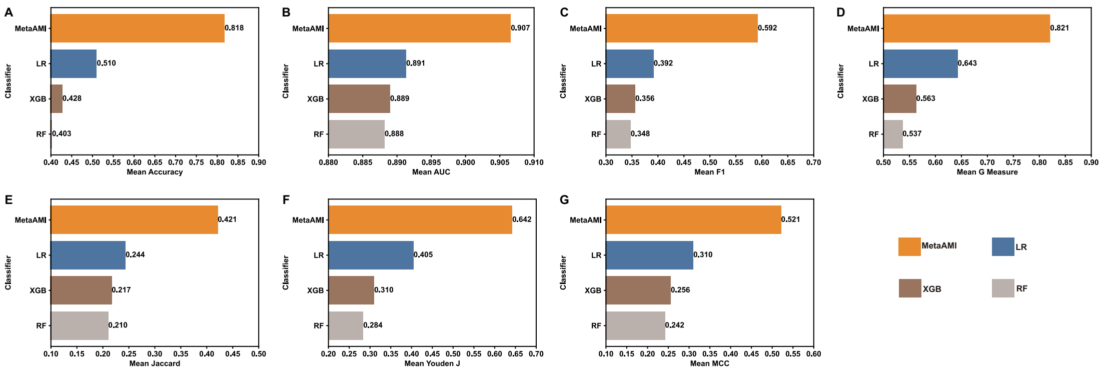


**Supplementary Fig. 2 Performance evaluation of MetaAMI.** MetaAMI outperformed the baseline models across multiple evaluation metrics, including **(A)** Accuracy, **(B)** Area Under the Curve (AUC), **(C)** F1 Score, **(D)** G-Measure, **(E)** Jaccard Index, **(F)** Youden J, and **(G)** Matthews Correlation Coefficient (MCC). LR: Logistic Regression; XGB: Extreme Gradient Boosting; RF: Random Forest.

**Supplementary Table. 1 ICD diagnosis codes for AMI.** ICD codes are standardized diagnostic codes used to classify diseases and health conditions in clinical practice and administrative records. The table includes the ICD-9 and ICD-10 diagnosis codes used for patient identification and cohort construction.

| | |
|---|---|
| ICD-9 | 41000,41001,41002,41010,41011,41012,41020,41021,41022, 41030,41031,41032,41040,41041,41042,41050,41051,41052, 41060,41061,41062,41070,41071,41072,41080,41081,41082, 41090,41091,41092 |
| ICD-10 | I210,I2101,I2102,I2109,I211,I2111,I2119,I212,I2121,I2129,I213, I214,I219I220,I221,I222,I228,I229 |

**Supplementary Table. 2 Hyperparameters of the baseline classifiers and the meta-learner.** The table summarizes the hyperparameters used for the baseline classifiers and the meta-learner in the proposed meta-learning framework for in-hospital mortality prediction.

| classifier | hyper parameter |
|---|---|
| LR | C = 0.01; solver = "lbfgs"; max_iter = 1000; class_weight = "balanced" |
| SVM | C = 0.003; kernel = "linear"; class_weight = "balanced"; probability = True |
| RF | n_estimators = 300; max_depth = 6; max_features = "sqrt"; class_weight = "balanced" |
| XGB | n_estimators = 100; max_depth = 3; learning_rate = 0.03; subsample = 0.8; colsample_bytree = 1; scale_pos_weight = 5.0 |
| MLP | epochs = 100; batch_size = 128; learning_rate = 0.01; dropout = 0.10; l2 = 1e-4; hidden_layers = [126, 64] |
| AE-MLP | ae_epochs = 50; ae_learning_rate = 5e-4; latent_dim = 16; ae_hidden_layers = [256, 128, 64]; epochs = 40; batch_size = 128; learning_rate = 0.01; dropout = 0.10; l2 = 1e-5; classifier_hidden = [64, 32] |
| OT-MLP | epochs = 100; batch_size = 128; learning_rate = 1e-3; dropout = 0.10; l2 = 1e-5; hidden_layers = [128, 64]; ot_eps = 5; ot_iters = 100 |
| Transformer | epochs = 30; batch_size = 64; learning_rate = 3e-4; dropout = 0.10; l2 = 1e-4; d_model = 128; n_heads = 4; n_layers = 2; ffn_multiplier = 2 |
| Meta-SVM | C = 0.01; gamma = 0.001; class_weight = "balanced" |
| Meta-MLP | Epochs = 50; batch_size = 32; learning_rate = 1e-4; hidden_layers = 4 |
| Meta-XGB | n_estimators = 300; max_depth = 3; learning_rate = 0.03; subsample = 0.8; n_jobs = 8 |
| Meta-LR | C = 0.01; solver = "lbfgs"; max_iter = 1000; class_weight = "balanced" |
| Meta-RF | n_estimators = 300; max_depth = 6; max_features = "sqrt"; class_weight = "balanced" |
| Meta-Transformer | epochs = 30; batch_size = 64; learning_rate = 3e-4; dropout = 0.10; l2 = 1e-4; d_model = 128; n_heads = 4; n_layers = 2; ffn_multiplier = 2 |

**Supplementary Table. 3 McNemar's test results of MetaAMI and baseline models.** For each comparison, the table reports the number of cases correctly classified by both models, correctly classified only by MetaAMI, correctly classified only by the baseline model, and misclassified by both models. A $p$ value < 0.05 was considered statistically significant.

| compare | Both correct | MetaAM correct Others wrong | MetaAM wrong Others correct | Both wrong | p_value | p_adj |
|---|---|---|---|---|---|---|
| MetaAMI vs LR | 2592 | 2196 | 153 | 913 | <0.0001 | <0.0001 |
| MetaAMI vs RF | 1937 | 2851 | 160 | 906 | <0.0001 | <0.0001 |
| MetaAMI vs XGB | 2068 | 2720 | 157 | 909 | <0.0001 | <0.0001 |
| MetaAMI vs AE-MLP | 4230 | 558 | 118 | 948 | 5.84E-64 | 2.92E-63 |
| MetaAMI vs MLP | 4255 | 533 | 131 | 935 | 1.32E-54 | 5.29E-54 |
| MetaAMI vs SVM | 4383 | 405 | 88 | 978 | 5.81E-46 | 1.74E-45 |
| MetaAMI vs OT-MLP | 4404 | 384 | 128 | 938 | 1.86E-29 | 3.71E-29 |
| MetaAMI vs Transformer | 4344 | 444 | 251 | 815 | 3.27E-13 | 3.27E-13 |

**Supplementary Table. 4 Features selected by expert.** The table lists the features selected by clinician from the features identified in the study workflow, based on their clinical relevance to AMI and in-hospital mortality prediction.

| | |
|---|---|
| age | history_of_falling_within_3_mnths |
| aids | inr_pt |
| anion_gap | lactate |
| arterial_o2_pressure | magnesium |
| base_excess | malignant_cancer |
| bicarbonate | mch |
| braden_activity | mchc |
| braden_mobility | mcv |
| braden_nutrition | metastatic_solid_tumor |
| cerebrovascular_disease | mild_liver_disease |
| charlson_comorbidity_index | non_invasive_blood_pressure_diastolic |
| chloride | non_invasive_blood_pressure_mean |
| chronic_pulmonary_disease | non_invasive_blood_pressure_systolic |
| congestive_heart_failure | paraplegia |
| creatine_kinase_mb_isoenzyme | pco2 |
| creatinine | peptic_ulcer_disease |
| dementia | peripheral_vascular_disease |
| diabetes_with_cc | ph |
| diabetes_without_cc | phosphate |
| gcs_eye_opening | platelet_count |
| gcs_motor_response | po2 |
| potassium | rdw |
| red_blood_cells | urea_nitrogen |
| renal_disease | white_blood_cells |
| respiratory_rate | gcs_verbal_response |
| rheumatic_disease | gender_binary |
| severe_liver_disease | glucose |
| sodium | heart_rate |
| stay_id | hematocrit |
| subject_id | hemoglobin |
| temperature_fahrenheit | pressure_ulcer_present |
| troponin_t | pt |
| ptt | |
